# Dopamine buffering capacity imaging: A pharmacodynamic fMRI method for staging Parkinson disease

**DOI:** 10.1101/817106

**Authors:** Kevin J. Black, Haley K. Acevedo, Jonathan M. Koller

## Abstract

We propose a novel brain imaging method for objectively quantifying disease severity in Parkinson disease (PD). Levodopa pharmacological fMRI (phMRI) hysteresis mapping is based on the clinical observation that the benefit from a dose of levodopa wears off more quickly as PD progresses. Biologically this has been thought to represent decreased buffering capacity for dopamine as nigrostriatal cells die. Buffering capacity has been modeled previously based on clinical effects, but clinical measurements are influenced by confounding factors such as patient fatigue. The new method proposes to measure the effect directly and objectively based on the timing of the known metabolic and blood flow response of several brain regions to exogenous levodopa. Such responses are robust and can be quantified without ionizing radiation using perfusion MRI.

Here we present simulation studies based on published clinical dose-response data and an intravenous levodopa infusion. Standard pharmacokinetic-pharmacodynamic methods were used to model the response. Then the effect site rate constant *k*_*e*_ was estimated from simulated response data plus Gaussian noise.

Predicted time:effect curves sampled at times consistent with phMRI differ substantially based on clinical severity. Estimated *k*_*e*_ from noisy input data was recovered with good accuracy.

These simulation results support the feasibility of levodopa phMRI hysteresis mapping to measure the severity of dopamine denervation objectively and simultaneously in several brain regions.

## Introduction

> *The intensity and duration of the effect after injection appear to correlate with the degree of akinesia, the action of L-DOPA lasting longer the less pronounced the akinesia.*
>
> — — Hirschmann and Mayer, 1964 (translated) (Hirschmann & Mayer, 1964)

Parkinson disease (PD) is characterized by progressive death of cells projecting from the substantia nigra to the striatum. One of the most important unmet needs in PD is to find objective, quantitative *in vivo* biomarkers of disease severity. Biomarkers of nigrostriatal denervation are sought for several important reasons, including as surrogate markers of disease progression in treatment trials (Gwinn et al., 2017; Ryman & Poston, 2019). Putative imaging biomarkers of disease progression include striatal [^18^F]fluorodopa PET or [^123^I]ioflupane SPECT. Unfortunately, these techniques do not accurately quantify nigrostriatal cell loss (Karimi et al., 2013). Presynaptic dopaminergic imaging of the midbrain does (Brown et al., 2013); nevertheless, alternative methods would be welcome.

Here we describe a novel potential biomarker, based on the clinical observation that the benefit from a dose of levodopa wears off more quickly as PD progresses. Early in the course of disease, a small dose of levodopa provides benefit long after the plasma levodopa concentration has declined substantially from its peak. The body responds as if the levodopa in the plasma filled a reservoir and then slowly leaked out to produce benefit. With disease progression, even though the same amount of levodopa circulates in the blood, the benefit wears off much faster, as if the reservoir had become leakier. Biologically, the reservoir may represent the diminishing buffering capacity of ascending dopaminergic axons as midbrain dopamine neurons die off (Marsden & Parkes, 1976). This wearing off of benefit has been quantified by a mathematical model that postulates a central effect compartment (reservoir) whose concentration of levodopa directly determines the clinical benefit. The buffering capacity in this model can be characterized by a single number, the effect site rate constant *k*_*e*_, which can be computed from serial measurements of both plasma concentration and clinical status (Sheiner, Stanski, Vozeh, Miller, & Ham, 1979). On average, patients with more severe PD and longer disease duration have a larger (“leakier”) *k*_*e*_ when modeled this way (Fig. 1) (M Contin et al., 2001, 1994; Harder & Baas, 1998; J G Nutt & Holford, 1996; J G Nutt, Woodward, Carter, & Gancher, 1992; Triggs et al., 1996). In fact, *k*_*e*_ can be the strongest predictor of the kinetics of response to levodopa in PD (M Contin et al., 2001; J G Nutt & Holford, 1996). Dopamine buffering capacity as measured by *k*_*e*_ also correlates significantly with nigrostriatal denervation as measured by DOPA uptake (Dietz et al., 2001) or dopamine transporter imaging (Manuela Contin et al., 2003). Unfortunately, the clinical measurements used to determine *k*_*e*_ are influenced by confounding factors such as patient fatigue and motivation, which likely add variance to the measurement. A direct, objective brain measure of response to LD may reduce this added variance.

**Figure 1.**
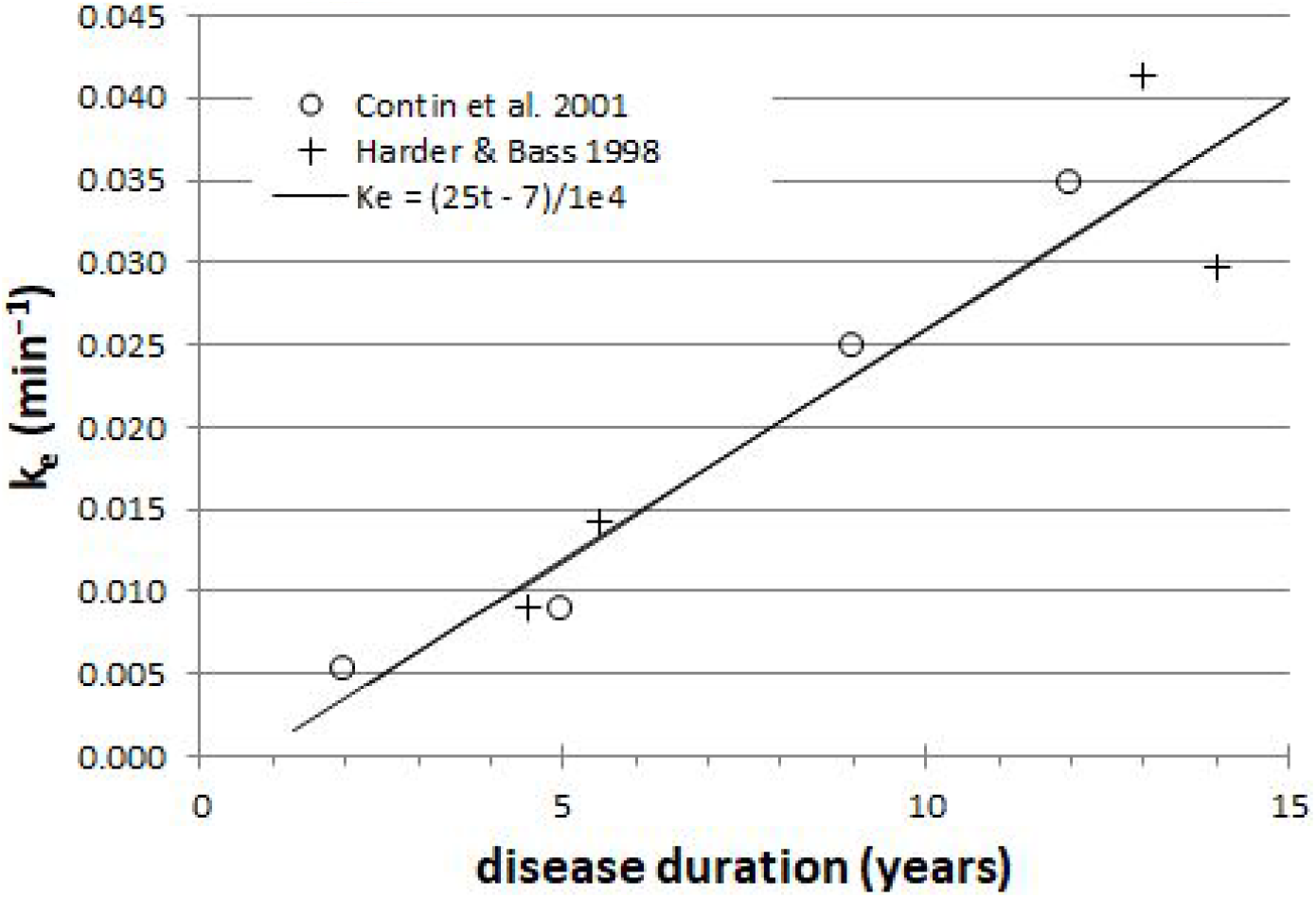
Across groups of PD patients, *k*_*e*_ is a surrogate for disease duration (*r*=0.95).

The effect of levodopa on the brain can be seen by measuring movement, but also by measuring regional cerebral blood flow (rCBF), reflecting regional brain activity (Black, Hershey, Hartlein, Carl, & Perlmutter, 2005; Hershey et al., 2003; Hershey, Black, Carl, & Perlmutter, 2000; Hershey et al., 1998). Crucially, using quantitative techniques, levodopa has no direct vascular effects after adequate carbidopa pretreatment (Hershey et al., 2003, 2000, 1998). Levodopa’s regional CBF effects reflect its regional effects on glucose metabolism and are prominent in pons and midbrain, thalamus, middle frontal gyrus, insula, putamen and cingulate cortex (Black et al., 2005; Hershey et al., 2000). Drug effects on rCBF in PD can be quantified without ionizing radiation using arterial spin labeling (ASL) perfusion MRI (Black, Koller, Campbell, Gusnard, & Bandak, 2010; Chen, Pressman, Simuni, Parrish, & Gitelman, 2015; Stewart, Koller, Campbell, & Black, 2014). The midbrain rCBF response to levodopa is robust whether measured with [^15^O]water PET (Black et al., 2005; Hershey et al., 2003, 2000, 1998) or with perfusion MRI (Chen et al., 2015).

Here we show, using simulated data based on published results in human PD patients, that quantifying dopamine buffering capacity *k*_*e*_ is likely to be feasible with existing technology.

## Methods

### Pharmacokinetics

Measuring *k*_*e*_ with levodopa phMRI would be infeasible if one had to repeatedly image a subject until a dose of levodopa wore off completely, perhaps for several hours in early PD. Fortunately, with faster wearing-off as PD progresses, there is also faster “wearing-on” or onset of drug effect (M Contin et al., 1990; John G Nutt, Carter, Lea, & Sexton, 2002; J G Nutt et al., 1992; Sohn et al., 1994). In fact, with a completely unrelated drug that also shows equilibration delay, giving the drug as a rapid intravenous (i.v.) infusion followed by a slow maintenance infusion allowed estimating the *k*_*e*_ just as precisely from the first 20 minutes of data as from 3½ hours of data (Sheiner et al., 1979). Fortunately we have used exactly this approach to dose levodopa in PD: a fast i.v. loading dose followed by a slow maintenance infusion (Black et al., 2003) (Fig. 2). This infusion method allows us to transiently achieve plasma levodopa concentrations of 1500-3500ng/mL, so we can measure *k*_*e*_ from both the rapid rise and fall of plasma levels.

**Figure 2.**
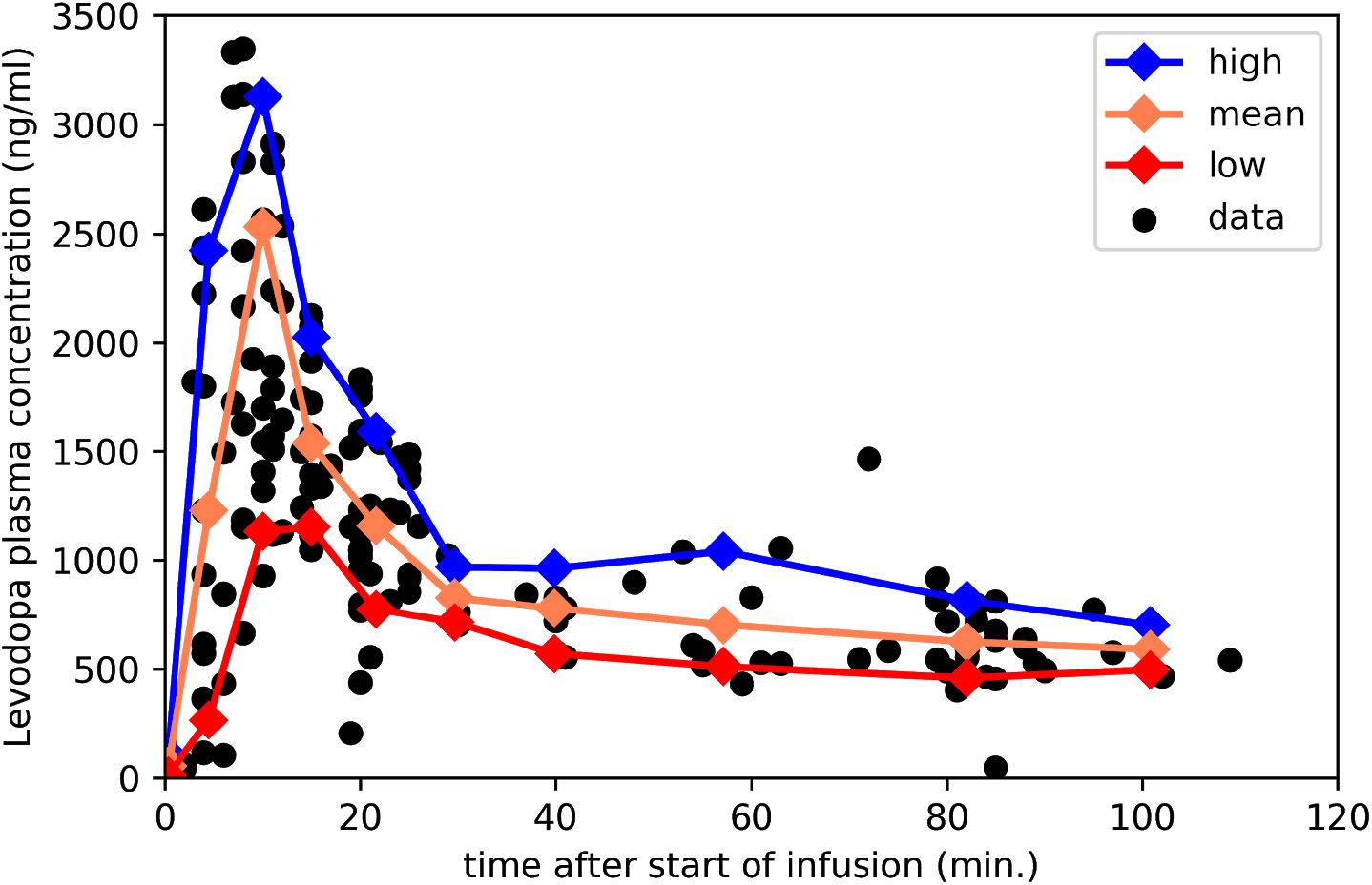
Plasma levodopa concentrations in PD patients following the “final dose” intravenous infusion method in ref. (Black et al., 2003). The 3 lines mark the mean, 90th and 10th percentile for samples collected in the corresponding intervals. Redrawn from data reported in ref. (Black et al., 2003).

For present purposes, likely time:concentration curves *C*_*p*_(*t*) in people with PD were taken from this infusion protocol, which aims to produce a steady-state levodopa concentration of 600ng/ml, and consists of a 10‑minute loading dose at 0.6426mg/kg followed by a maintenance infusion at 2.882 × 10^−5^ mg/kg/min × (140 yr – age)/yr (Black et al., 2003). For a 65-year-old 70-kg person that means 45mg over 10 min followed by 0.15mg/min, for a total dose over 150 min of 66mg. That i.v. dose is bioequivalent to 78mg oral levodopa (Robertson et al., 1989), though of course i.v. dosing leads to much higher transient peak plasma concentrations.

Using those data, we aggregated individual data points by time bins and plotted the mean, to estimate the most likely *C*_*p*_*(t)*, and the 10th and 90th percentile, to deal with a range of metabolic rates in patients (Fig. 2). The 10th percentile curve looked biologically unlikely, and for the simulations below was replaced by 70% of the mean curve, which produced similar predicted effect curves (see Jupyter notebook in supplementary material). The NumPy and matplotlib libraries in Python were used for simulations and data visualization (Hunter, 2007; Oliphant, 2006).

### Modeling the effect compartment

Holford and Sheiner describe the theoretical background for the effect compartment model (N. H. Holford & Sheiner, 1981). A later paper by Sheiner’s group simplifies the modeling with the assumption that *C*_*e*_ = *C*_*p*_ at steady state, leading to the definition of the effect compartment concentration curve by the simpler differential equation *C*_*e*_′ = *k*_*e*_(*C*_*p*_ − *C*_*e*_) (Unadkat, Bartha, & Sheiner, 1986).

If we use piecewise linear interpolation to estimate *C*_*p*_(*t*) between blood samples (as did ref. (Unadkat et al., 1986)), *C*_*e*_ can be computed in closed form. We can write *C*_*p*_ as *C*_*p*_(*t*−*t*_*i*_) = *C*_*p*_(*t*_*i*_) + *m*_*i*_(*t*−*t*_*i*_) on the interval [*t*_*i*_, *t*_*i*+1_], where *m*_*i*_ = [*C*_*p*_(*t*_*i*+1_)−*C*_*p*_(*t*_*i*_)] / [*t*_*i*+1_−*t*_*i*_]. We need a value for the effect site concentration before the infusion starts, *C*_*e*_(*t*_0_). For the purposes of this report, we can reasonably assume *C*_*e*_(*t*_0_) = *C*_*p*_(*t*_0_), which will be approximately true if at the time of the first blood draw patients have refrained from taking oral levodopa for 8-10 hours, since *t*_½*e*_ is <5 hours, and usually <2.5 hours (see Table 4 in ref. (M Contin et al., 2001)).

The solution to this initial value problem is

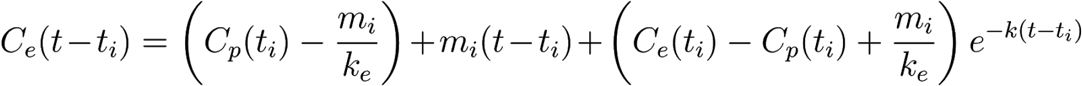

defined on the interval (*t*_*i*_, *t*_*i*+1_] (Maxima, 2014).

### Predicting effect from levodopa concentrations in PD

To test this method, one needs to estimate a reasonable variety of time:effect curves in PD. Not only *k*_*e*_ but also the concentration–effect curve changes with disease severity. Contin and colleagues showed that a sigmoid *E*_*max*_ model reasonably fit the data from a wide range of PD severity (M Contin et al., 2001). We adopt their measurements of *EC*_50_, *n* (the Hill coefficient) and *k*_*e*_ for a variety of disease severity groups; namely, means for Hoehn and Yahr (Hoehn & Yahr, 1967) stages I through IV in addition to the mildest and most severely affected individual subjects in the Contin *et al.* report (their Table 4). These parameters and the sigmoid *E*_*max*_ model are combined to create time–effect curves that we could expect from a brain region whose activity changes reliably with increased dopamine release in the brain with administration of exogenous levodopa. Note that the dopa-responsive region need not itself contain dopamine receptors, but rather may be a “downstream” region (*e.g.* motor cortex) innervated by cells that do (*e.g.* posterior putamen). The simulated data use a baseline CBF of 50 ml/hg/min and maximal effect was set at 35 ml/hg/min, consistent with a ~70% rCBF increase in midbrain after a relatively large levodopa-carbidopa dose (Chen et al., 2015).

### Adding noise

The brain imaging time:effect curves assessed by any real brain imaging method will not be perfect, noise-free estimates, but will be contaminated by variability from biological or instrumentation issues. To test how well we can expect to recover *k*_*e*_ (and the other pharmacodynamic parameters) from a real experiment, we add noise to the simulated data described in the previous paragraph. We added Gaussian noise with a coefficient of variation (CoV) of 12.9%; this value was chosen based on the CoV in a cortical gray matter region across sixteen 34-second CBF images in 11 adults with PD scanned with a pCASL sequence while fixating a crosshair (unpublished data, K. J. Black and colleagues).

### Parameter estimation

We simultaneously estimated k_e_, EC_50_, and n from the data, given the model, using the `lmfit` package in Python (`ampgo` followed by `emcee` modules) (Newville et al., 2019).

### Accuracy

The accuracy of the method was tested by comparing the estimated *k*_*e*_ to the input *k*_*e*_. Secondary similar analyses were done for *EC*_50_ and *n*.

## Results

### Predicted LD time:concentration curves in effect compartment

Fig. 3 shows the expected concentration over time in the effect compartment, depending on the severity of PD. One can easily appreciate the faster exchange between plasma and the effect compartment when *k*_*e*_ is high (*i.e.*, when the equilibration half-life *t*_½*e*_ is short).

**Figure 3.**
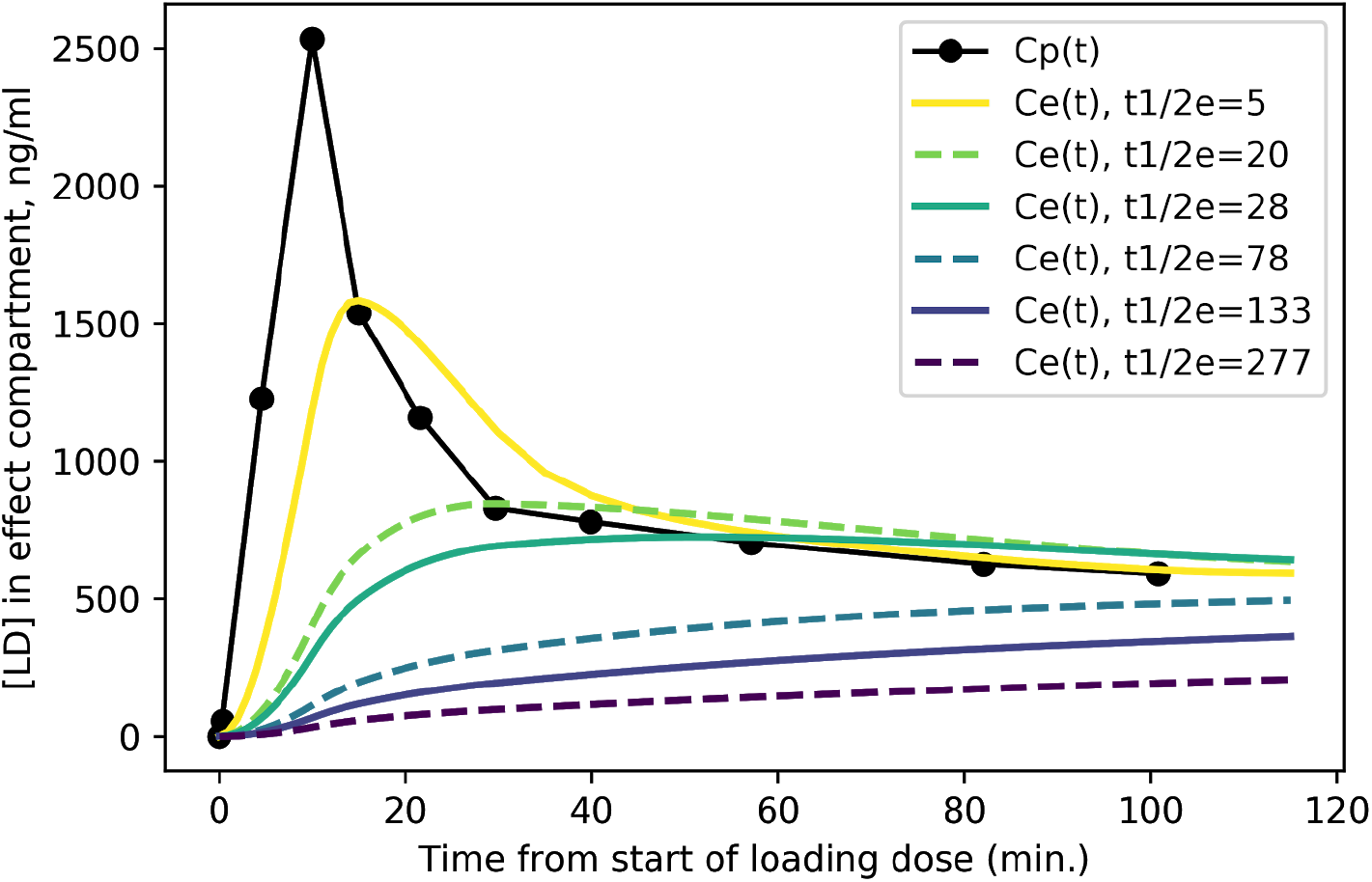
Predicted levodopa concentration in effect compartment at various disease severity levels. Curves are labeled by *t*_½*e*_ = *ln* 2/*k*_*e*_ from more severe PD (*t*_½*e*_ = 5 min.) to milder PD (*t*_½*e*_ = 277 min.).

### Time:effect curves by disease severity

We modeled the expected rCBF response in midbrain to the rapid i.v. infusion, based on published levodopa pharmacokinetics in PD (Black et al., 2003) and published mean pharmacodynamic parameters for Hoehn & Yahr stages I, II, III, and IV (M Contin et al., 2001; Harder & Baas, 1998). The predicted signals are quite distinct, assuming a typical *C*_*p*_(*t*) time:plasma curve (Fig. 4A). If a given subject’s pharmacokinetics produce higher plasma levels, the distinctions are still fairly clear (Fig. 4B). Of course, if an individual’s plasma levels are low, an effect may not be evident, especially in more severe PD (Fig. 4C).

**Figure 4.**
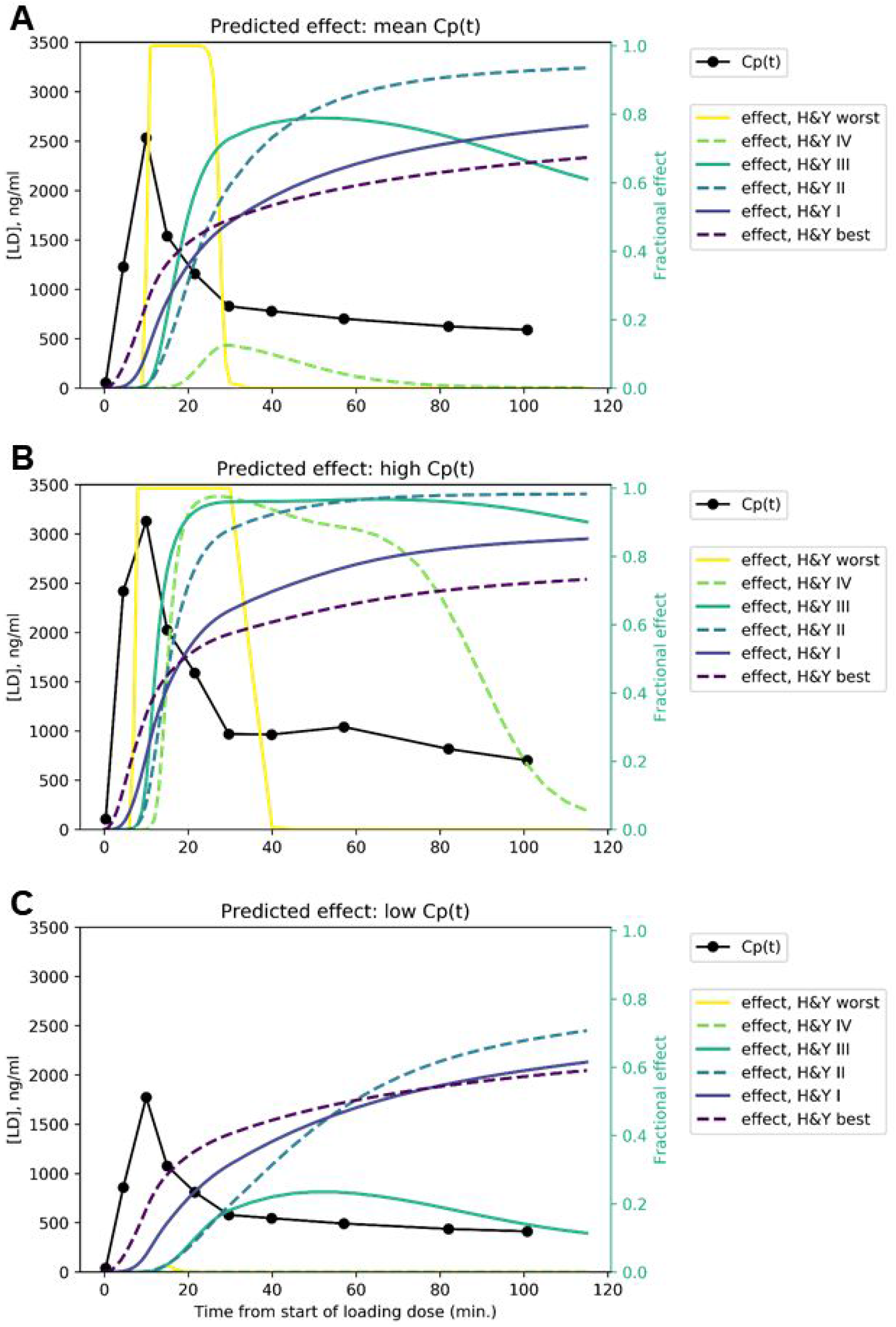
Predicted time:effect curves at various disease severity levels assuming **A)** mean, **B)** high and **C)** low *C*_*p*_(*t*) in response to the levodopa infusion.

### Accuracy

*k*_*e*_ estimated from time:effect curves in the presence of noise was generally accurate (Fig. 5; see also Table S1). More advanced disease led to more distinct predicted time–activity curves (see Fig. 4A), reflected in more accurate results (Fig. 5A-C).

**Figure 5.**
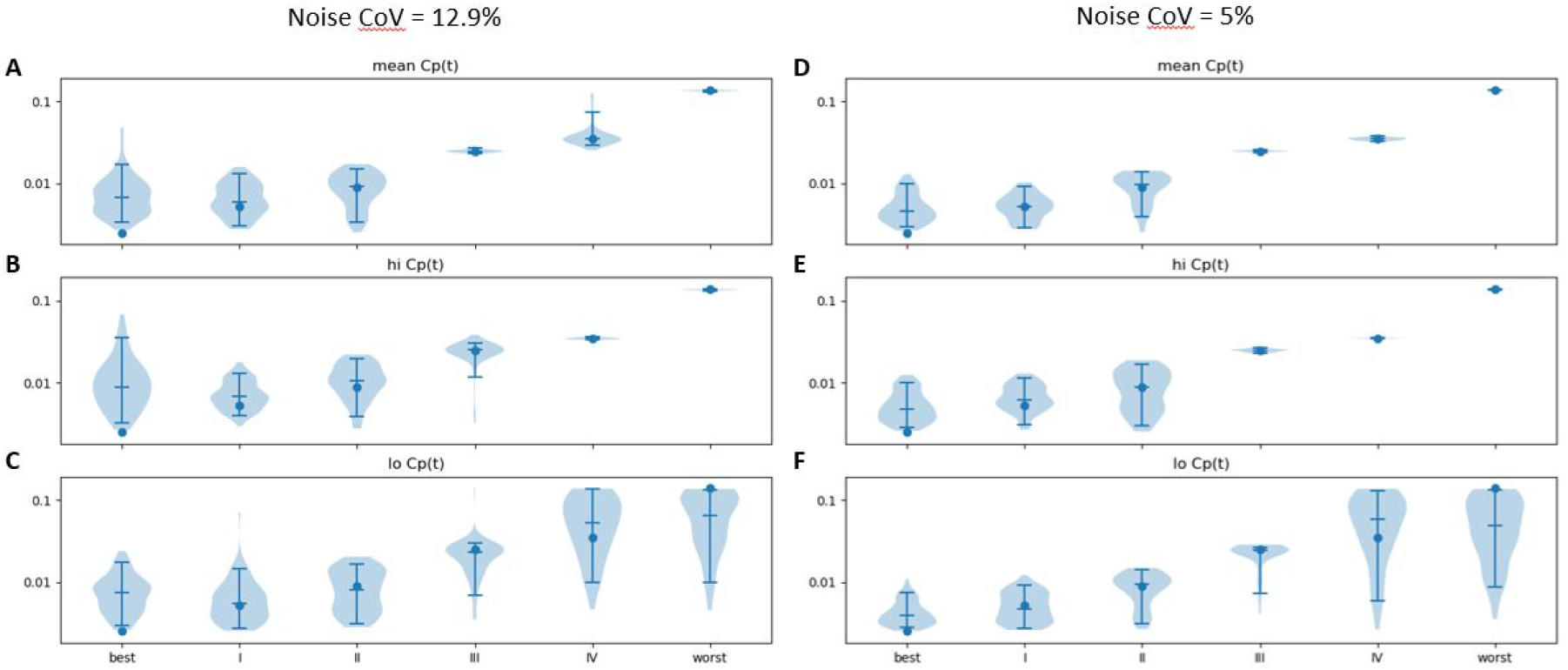
Estimated *k*_*e*_ (vertical axis) across 100 sets of noise added to the time:effect curve computed for the *k*_*e*_ (horizontal axis), *EC*_50_, and *n* for various severities of PD as reported in ref. (M Contin et al., 2001), assuming **A)** mean, **B)** high and **C)** low *C*_*p*_(*t*) in response to the levodopa infusion. Noise CoV = 12.9%. Width of plot is proportional to frequency of output of the given magnitude. Filled circle: input n. Horizontal lines note the 5th, 50th and 95th percentiles. At right, similar results are shown for noise CoV = 5% for **D)** mean, **E)** high and **C)** low *C*_*p*_(*t*) responses to the LD infusion.

Results were more accurate if the noise was reduced from a CoV of 12.9% to 5% (Fig. 5D-F; see also Table S2). Similar plots for *EC*_50_ and *n* are provided as Supplementary Information.

In an attempt to improve further the accuracy, we examined the effect of spreading the LD infusion over twice the time. We hypothesized that the limited temporal resolution of the perfusion MR method, combined with the relatively small timing difference in onset of action in mild *vs*. very mild PD, limited discrimination at the milder end of the severity range. Results are shown in Fig. 6.

**Figure 6.**
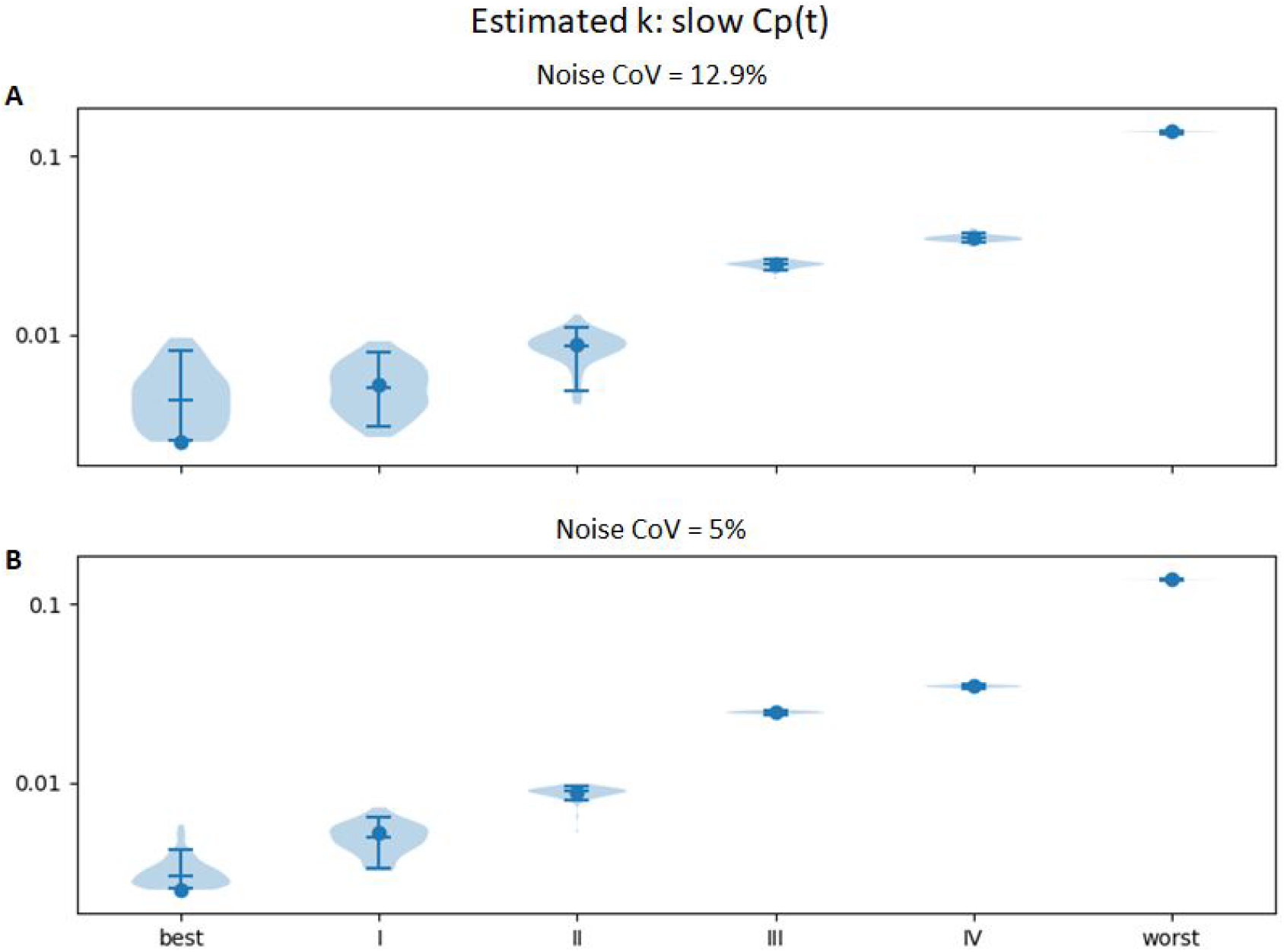
Same Estimated *k*_*e*_ (vertical axis) across 100 sets of noise added to the time:effect curve computed for the *k*_*e*_ (horizontal axis), *EC*_50_, and *n* for various severities of PD as reported in ref. (M Contin et al., 2001), with *C*_*p*_(*t*) estimated for an levodopa infusion twice as long (at half the rate, so that the total infused dose is equivalent). **A)** noise CoV = 12.9%; **B)** noise CoV = 5%.

Similar plots for *EC*_50_ and *n* are provided as Supplementary Figures 5 and 6.

## Discussion

We present a novel brain imaging method for objectively quantifying disease severity in Parkinson disease (PD), which we refer to as dopamine buffering capacity imaging, or more precisely, levodopa phMRI hysteresis mapping. The temporally distinct time:effect curves predicted in Fig. 4 suggest that even with some imperfection in the rCBF signal, we can expect to derive a reasonably accurate *k*_*e*_ for a brain region that responds to exogenous levodopa with a clear dose-response curve.

Demonstrating efficacy for potential disease-modifying therapies in PD has been difficult. Delayed start designs and similar approaches that rely on change in clinical severity over time require years to complete, large patient groups, and even then have not yet been successful (Hurley, 2011). A validated biomarker would be of great value in improving this situation. The Michael J. Fox Foundation for Parkinson’s Research designates “the identification, development and use of biomarkers to diagnose and track Parkinson’s disease” as a priority area, noting that a successful biomarker “would mean better disease management for patients” and “improve and speed clinical development of disease-modifying therapies” (The Michael J. Fox Foundation for Parkinson’s Research, n.d.). In simulated data based on published results and reasonable assumptions, levodopa phMRI hysteresis mapping appears likely to fill that need.

Of course one does not need an MRI machine to tell if a drug is improving movement in PD, and the present proposal draws on previous studies using pharmacokinetic-pharmacodynamic (PK-PD) modeling of tapping speed or UPDRS score response to levodopa challenge. However, the assessment of drug response using brain imaging is novel, and provides several potential advantages. The rCBF response is objective, rater-independent, and does not require subject movement. Furthermore, buffering capacity is measured simultaneously in numerous brain regions rather than just the motor system, potentially informing pathophysiological research on the increasingly recognized nonmotor symptoms of PD (Kehagia, 2016).

Chan, Nutt and Holford have subsequently extended the PK-PD model with the aim of better modeling long-term changes with disease progression in PD (Chan, Nutt, & Holford, 2005; N. H. G. Holford et al., 2006). Their revised model includes factors intended to account for clinical observations like morning benefit and the long-duration response, and in their data *k*_*e*_ (reported as *T*_*eqf*_ = *ln* 2 / *k*_*e*_) did not change significantly over time. However, as they note, other factors could explain the difference in results, and their more complicated model was made possible by a very large set of longitudinal data. While the extended model may be ideal for optimal understanding of physiology from clinical PK-PD data, it is not essential for the present purpose of identifying a biomarker of nigrostriatal denervation in PD. In other words, if *k*_*e*_ as derived from the model we use correlates highly with disease severity, it will serve its intended purpose just fine.

Every step of this method has been proven individually: i.v. levodopa has been used safely for over 50 years (Siddiqi et al., 2015); the infusion method described for the simulated data has been used in over 100 subjects (refs. (Black et al., 2003, 2010) and Black et al., unpublished data); levodopa concentration can be quantified accurately in plasma (Karimi, Carl, Loftin, & Perlmutter, 2006); the response to levodopa can be measured by ASL fMRI (Black et al., 2010; Chen et al., 2015; Stewart et al., 2014); midbrain has a robust rCBF response to single, clinically sensible doses of levodopa (Black et al., 2005; Chen et al., 2015; Hershey et al., 2003, 2000, 1998), and software exists for estimating PK-PD parameters from fMRI data on a voxel-by-voxel or regional level (Koller, Vachon, Bretthorst, & Black, 2016; Newville et al., 2019). In other words, every part of the method described here is well proven; it is their combination and interpretation as a disease severity measure that is novel.

### Foreseeable obstacles and possible solutions

Some potential difficulties in implementing dopamine buffering capacity imaging are foreseeable, but can be mitigated. These include a need for high temporal resolution, uncertain optimal dosing, head movement during MRI, and variable attention and alertness during the scans.

#### Temporal resolution

Prior data showing robust rCBF responses to levodopa averaged data across a group and over several scans in the pre- and on-drug conditions, *i.e.* with a time resolution of about 30 minutes. Measuring dopamine buffering capacity in individual subjects pushes the envelope, requiring measuring response to levodopa in single subjects and at a time resolution of 1-2 min or better. Fortunately, current pCASL methods allow an unbiased whole-brain measure of blood flow in about 5-35 seconds. However, these images are statistically noisy. If estimated *k*_*e*_ proves less accurate with individual subject data than these simulations predict, additional information contained in the data may strengthen prediction of disease severity. Specifically, from the plasma levodopa concentration curve and the MRI response data one computes not only *k*_*e*_ but also *EC*_50_ and *n*, which also change with disease severity (M Contin et al., 2001). Possibly combining all three parameter estimates may more accurately measure disease severity.

#### Optimal dosing

Subjects with more advanced disease will show little response if they also happen to have low plasma levodopa levels. Solutions could include higher dosing for more severely affected individuals, though this choice could increase the risk of dyskinesias in the scanner that could affect comfort or head movement. Alternatively, if needed, one could estimate the optimal dose for each subject with, say, a single small test dose of i.v. levodopa with a pre- and post-drug blood sample, on a day prior to the scan day.

#### Head movement

In our experience, most PD patients do well holding the head still during an MRI session. However, acquiring a single CBF image can take 6-34 s (Wang et al., 2012), an interval long enough that head movement on the scale of mm may be nontrivial. Participants with levodopa-induced dyskinesias may have additional head movement. Within-frame head movement adds to variance and may bias quantitative estimates. Solutions may include more rigid head fixation, shorter repetition times (TRs), prospective motion correction, or removing or underweighting CBF images compromised by movement (Tanenbaum, Snyder, Brier, & Ances, 2015).

#### Attention / alertness

In initial pilot studies, we find that several factors combine to make continued alertness throughout the scan period difficult: PD patients often have insomnia, the scans are long and repetitive, and levodopa contributes to sleepiness. Solutions may include adding an attention task (though that will change resting brain activity), study staff repeatedly awakening the participant, or monitoring for alertness and removing or accounting statistically for frames during which the participant appears asleep.

### Next steps

The simplest first step to validating this method is correlative in nature. Specifically, one would enroll people with a wide variety of PD severity and compare regional *k*_*e*_ values, most likely in midbrain or posterior putamen, to clinical measures of disease severity such as off-period UPDRS scores (Fahn, Elton, & UPDRS program members, 1987). More definitive validation of dopamine buffering imaging may include longer-term or autopsy studies in patients, necropsy studies in animals with graded nigrostriatal lesions, or comparison to the recently validated midbrain [^11^C]DTBZ PET approach (Brown et al., 2013). If these studies are successful, the dopamine buffering capacity imaging method will beg for further application as a surrogate marker of disease severity in PD.

## Data accessibility statement

Code available at: https://bitbucket.org/kbmd/hysteresis/

## Authors’ contributions

KJB designed the study, generated the test data, and wrote the initial draft. HKA and JMK performed simulation testing and revised the text. All authors gave final approval for publication.

## Supplementary Materials

All of the figures and tables below are based on 100 iterations of added Gaussian noise.

**Figure S1.**
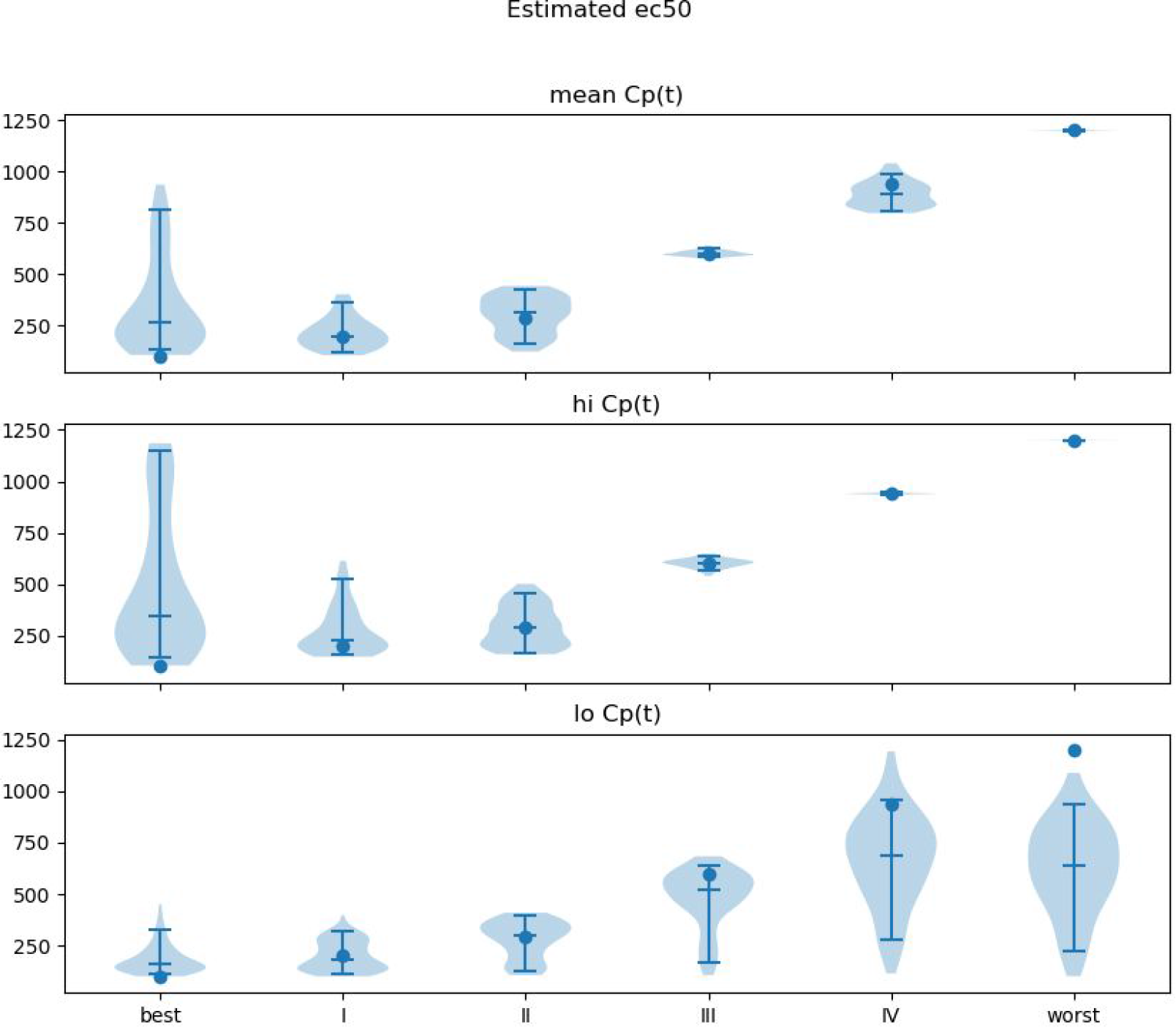
Accuracy of EC_50_, noiseCoV = 5%. Width of plot is proportional to frequency of output of the given magnitude. Filled circle: input EC_50_. Horizontal lines note the 5th, 50th and 95th percentiles.

**Figure S2.**
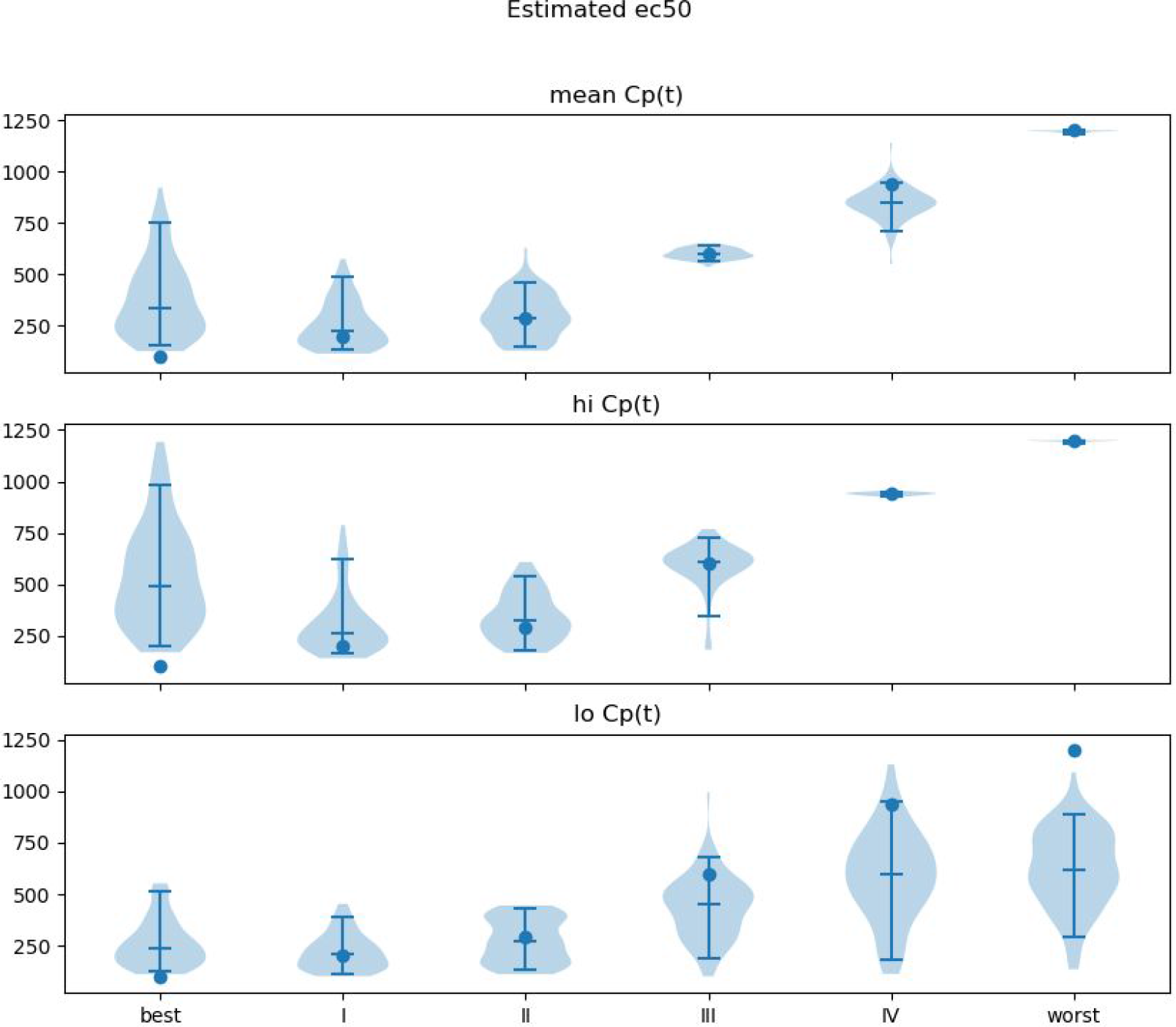
Accuracy of EC50, noiseCoV = 12.9%. Width of plot is proportional to frequency of output of the given magnitude. Filled circle: input EC_50_. Horizontal lines note the 5th, 50th and 95th percentiles.

**Figure S3.**
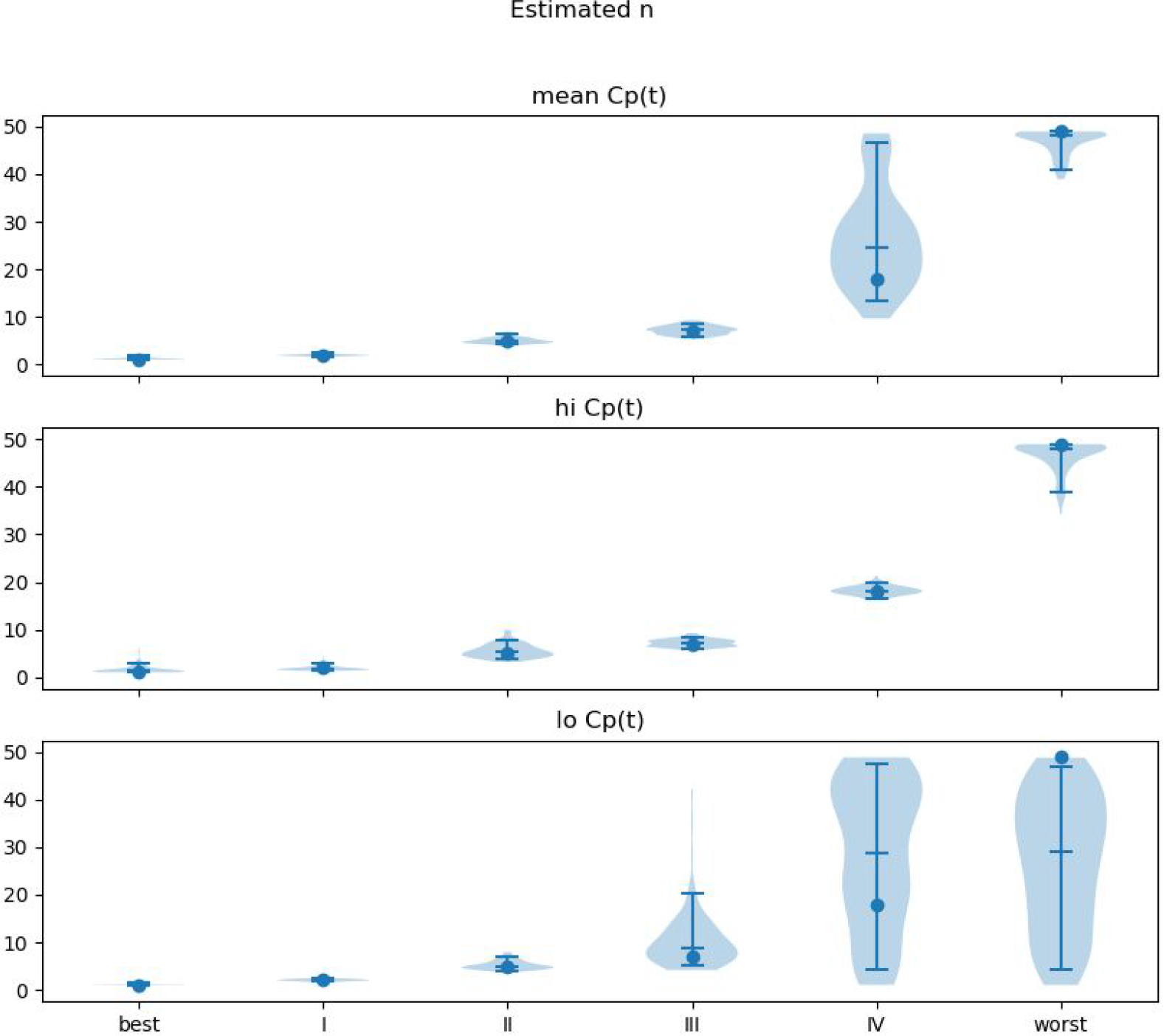
Accuracy of N, noiseCoV = 5%. Width of plot is proportional to frequency of output of the given magnitude. Filled circle: input *n*. Horizontal lines note the 5th, 50th and 95th percentiles.

**Figure S4.**
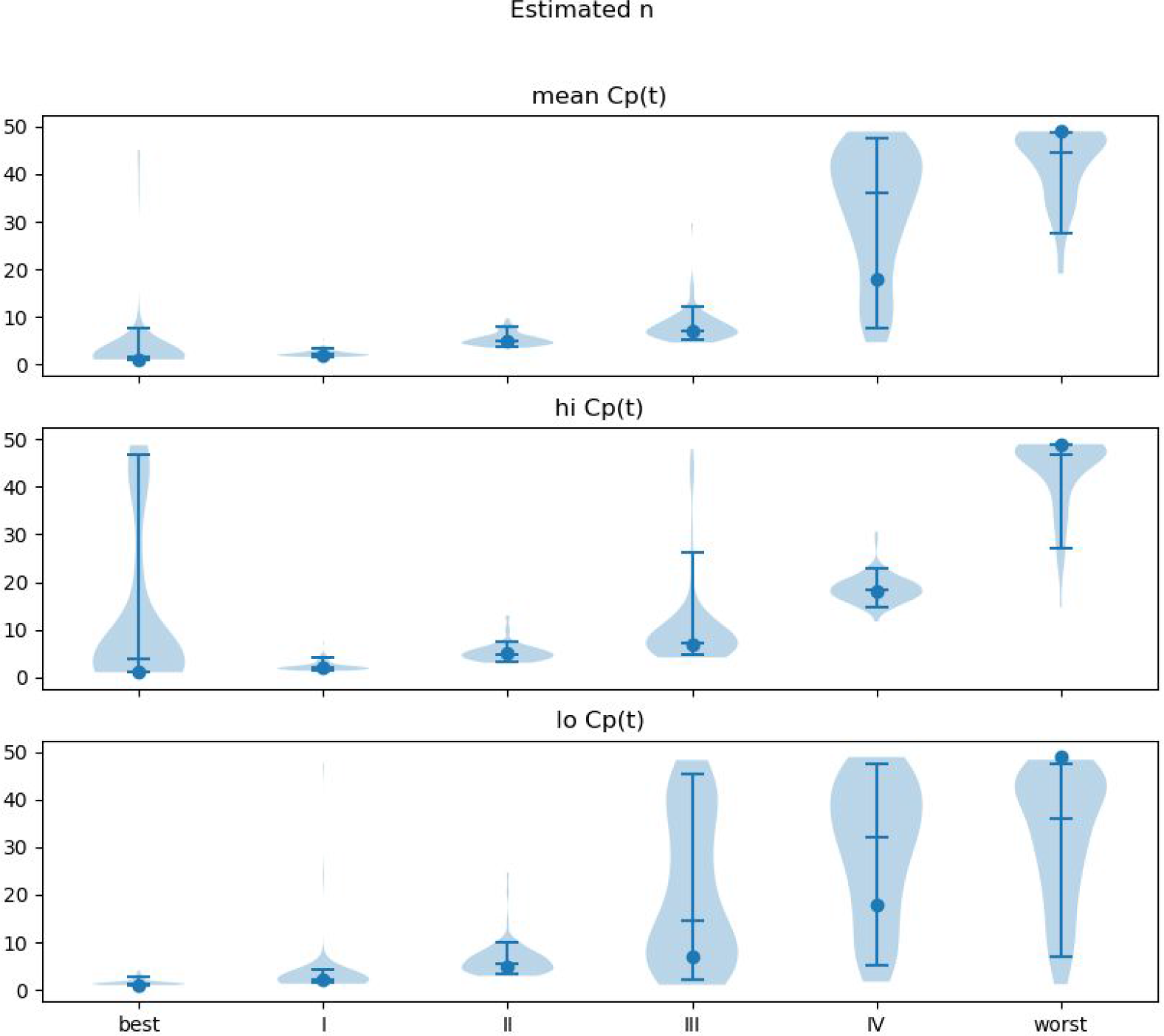
Accuracy of N, noiseCoV = 12.9%. Width of plot is proportional to frequency of output of the given magnitude. Filled circle: input *n*. Horizontal lines note the 5th, 50th and 95th percentiles.

**Figure S5.**
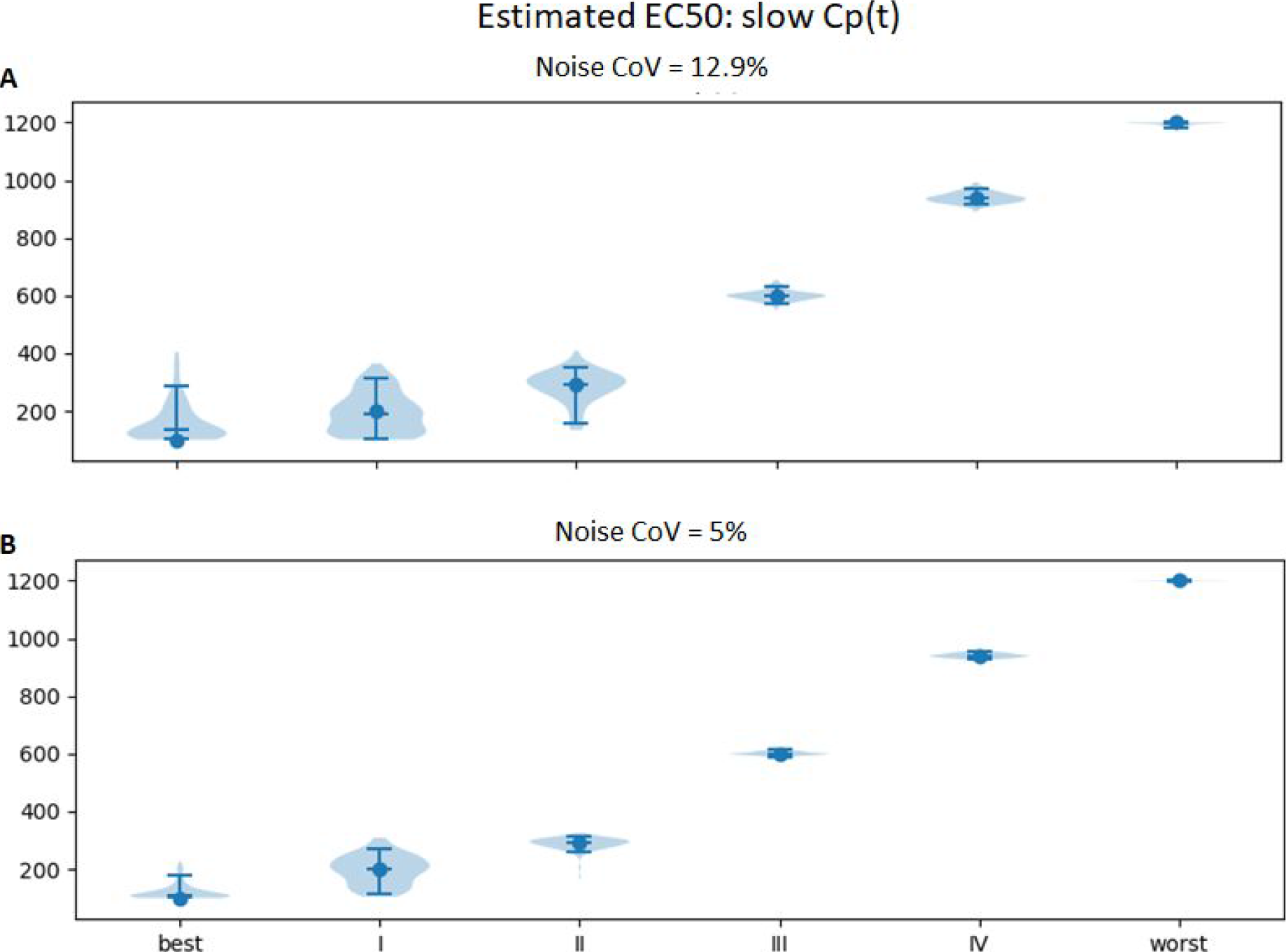
Accuracy of EC_50_, “slow” concentration curve. Width of plot is proportional to frequency of output of the given magnitude. Filled circle: input EC_50_. Horizontal lines note the 5th, 50th and 95th percentiles.

**Figure S6.**
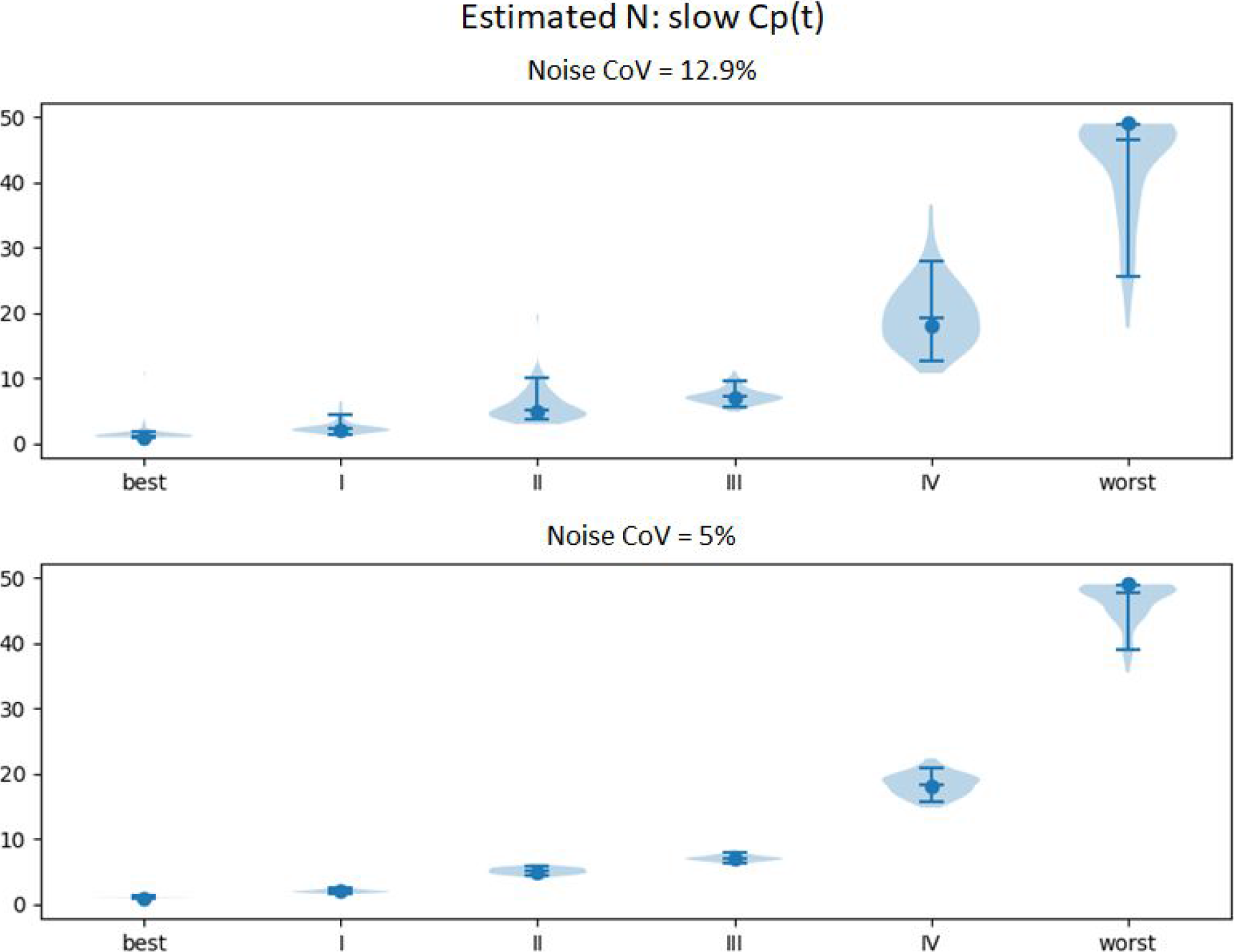
Accuracy of N, “slow” concentration curve. Width of plot is proportional to frequency of output of the given magnitude. Filled circle: input EC_50_. Horizontal lines note the 5th, 50th and 95th percentiles.

**Table S1.** Accuracy of ke, noise CoV = 12.9%. “Mean,” “hi,” and “lo” are the three time–activity curves described in the Pharmacokinetics section of Methods. Roman numerals refer to the mean pharmacodynamic values from the Hoehn and Yahr categories in ref. (M Contin et al., 2001). “Best” and “worst” refer to the extreme pharmacodynamic values from the same reference.

| **H&Y** | **Cp** | **target** | **median** | **5th %tile** | **95th %tile** |
| --- | --- | --- | --- | --- | --- |
| best | mean | 0.0025 | 0.0068 | 0.0034 | 0.0173 |
| I | mean | 0.0052 | 0.0059 | 0.0030 | 0.0131 |
| II | mean | 0.0089 | 0.0091 | 0.0033 | 0.0152 |
| III | mean | 0.0248 | 0.0247 | 0.0231 | 0.0269 |
| IV | mean | 0.0347 | 0.0349 | 0.0291 | 0.0747 |
| worst | mean | 0.1386 | 0.1374 | 0.1317 | 0.1385 |
| best | hi | 0.0025 | 0.0089 | 0.0033 | 0.0358 |
| I | hi | 0.0052 | 0.0069 | 0.0040 | 0.0129 |
| II | hi | 0.0089 | 0.0105 | 0.0038 | 0.0196 |
| III | hi | 0.0248 | 0.0251 | 0.0119 | 0.0306 |
| IV | hi | 0.0347 | 0.0346 | 0.0334 | 0.0363 |
| worst | hi | 0.1386 | 0.1381 | 0.1321 | 0.1386 |
| best | lo | 0.0025 | 0.0074 | 0.0029 | 0.0175 |
| I | lo | 0.0052 | 0.0054 | 0.0027 | 0.0146 |
| II | lo | 0.0089 | 0.0081 | 0.0031 | 0.0167 |
| III | lo | 0.0248 | 0.0230 | 0.0069 | 0.0296 |
| IV | lo | 0.0347 | 0.0529 | 0.0099 | 0.1357 |
| worst | lo | 0.1386 | 0.0656 | 0.0099 | 0.1333 |

**Table S2.** Accuracy of ke, noise CoV = 5%. “Mean,” “hi,” and “lo” are the three time–activity curves described in the Pharmacokinetics section of Methods. Roman numerals refer to the mean pharmacodynamic values from the Hoehn and Yahr categories in ref. (M Contin et al., 2001). “Best” and “worst” refer to the extreme pharmacodynamic values from the same reference.

| H&Y | Cp | target | median | 5th %tile | 95th %tile |
| --- | --- | --- | --- | --- | --- |
| best | mean | 0.0025 | 0.0046 | 0.0029 | 0.0100 |
| I | mean | 0.0052 | 0.0052 | 0.0029 | 0.0093 |
| II | mean | 0.0089 | 0.0097 | 0.0039 | 0.0137 |
| III | mean | 0.0248 | 0.0248 | 0.0241 | 0.0256 |
| IV | mean | 0.0347 | 0.0350 | 0.0325 | 0.0380 |
| worst | mean | 0.1386 | 0.1385 | 0.1364 | 0.1386 |
| best | hi | 0.0025 | 0.0047 | 0.0029 | 0.0101 |
| I | hi | 0.0052 | 0.0061 | 0.0030 | 0.0115 |
| II | hi | 0.0089 | 0.0087 | 0.0030 | 0.0168 |
| III | hi | 0.0248 | 0.0249 | 0.0232 | 0.0265 |
| IV | hi | 0.0347 | 0.0348 | 0.0343 | 0.0353 |
| worst | hi | 0.1386 | 0.1385 | 0.1363 | 0.1386 |
| best | lo | 0.0025 | 0.0039 | 0.0028 | 0.0074 |
| I | lo | 0.0052 | 0.0047 | 0.0028 | 0.0092 |
| II | lo | 0.0089 | 0.0093 | 0.0031 | 0.0140 |
| III | lo | 0.0248 | 0.0245 | 0.0073 | 0.0264 |
| IV | lo | 0.0347 | 0.0587 | 0.0059 | 0.1293 |
| worst | lo | 0.1386 | 0.0492 | 0.0086 | 0.1346 |

